# Inflammatory chemokine receptors CCR1, CCR2, CCR3 and CCR5 are essential for an optimal T cell response to influenza

**DOI:** 10.1101/2025.03.20.644330

**Authors:** Marieke Pingen, Catherine E. Hughes, Laura Medina-Ruiz, Heather Mathie, Jennifer A. Barrie, Chris AH Hansell, Robin Bartolini, Megan KL MacLeod, Gerard J Graham

## Abstract

Inflammatory chemokine receptors CCR1/2/3/5 (iCCRs) play an important role in the recruitment of immune cells involved in innate immune functions and orchestrating the adaptive immune response. Here we utilise an influenza A virus (IAV) challenge to investigate the combinatorial roles of the iCCRs in the anti-IAV immune response.

We did not observe any gross differences in infection-driven pathology in the absence of iCCRs. Despite iCCR deletion resulting in decreased migration of monocytes, migratory macrophages and B cells to lungs during acute IAV infection, no differences in dendritic cell numbers were observed. Whilst the total number of T cells was similar in lungs of iCCR-deficient mice, the number of IAV-specific CD4 but not CD8 T cells in the lung was strongly reduced in the absence of iCCRs. Furthermore, fewer CD4, but not CD8, T cells produced IFN-γ.

This CD4 T cell phenotype persisted into the memory stage of infection, with fewer IAV-specific and IFN-γ^+^ CD4 but not CD8 T cells at 29 days post infection.

In conclusion, despite having no impact on dendritic cell migration between the lung and the draining lymph node, iCCR deletion is associated with an altered CD4 T cell response to IAV infection.

## INTRODUCTION

The lung is constantly exposed to environmental challenges, including particles and pathogens, and a timely, balanced local immune response is critical to protect respiratory function^1^. As such there is a wide variety of innate and adaptive immune cells that reside in the lung to maintain homeostasis and provide the first line of defence against infection or damage^2^. Subsequently, when required, additional immune and inflammatory cells are recruited to the lung^3^.

Recruitment of immune cells to homeostatic and inflamed, or infected, tissues is predominantly regulated by chemokines, which are defined on the basis of a conserved cysteine motif and divided into CC, CXC, XC and CX3C subfamilies^4^. Upon inflammatory or immune challenge, both tissue-resident immune cells and structural cells can produce chemokines resulting in recruitment of additional immune and inflammatory cells expressing cognate chemokine receptors^4-6^. We have a particular interest in the inflammatory chemokine receptors CCR1, 2, 3 and 5 (hereafter referred to as iCCRs), which are expressed from a tight, highly conserved chromosomal locus^7^, and which predominantly orchestrate recruitment and intra-tissue movement of non-neutrophilic myeloid inflammatory cells^8, 9^. In addition to regulating inflammatory cell recruitment, these iCCRs and their ligands play important roles in orchestrating the adaptive immune response such as through regulation of antigen presenting cell activity and direct control of lymphocyte movement^4^. In particular, T cells are known to express iCCRs^4^ but their role in this cellular context is currently unclear. To study the expression of, and roles for, the iCCRs in immune and inflammatory responses we have generated 2 novel mouse strains. In one of these, iCCR KO mice, we have deleted the entire chromosomal locus incorporating these 4 receptors to assess their combinatorial contribution to the inflammatory response^8^. The other strain is a multi-receptor reporter mouse strain, REP mice, in which a BAC covering the iCCR locus has been engineered to replace the 4 receptor coding sequences with spectrally distinct fluorescent reporters. This BAC has then been transgenically inserted into a gene-free region ensuring that normal expression of the iCCRs is unaffected^9^.

In previous studies we used these mouse strains to determine the roles for iCCRs in controlling migration of innate immune cells, in particular myeloid cells such as monocytes and macrophages, using sterile models of inflammation. These studies have revealed a hitherto underappreciated specificity in the expression of iCCRs by leukocytes and their *in vivo* roles during migration^8, 9^. We have now turned our attention to the roles for iCCRs in the immune and inflammatory response to pathogen infection using a model of influenza A (IAV) infection. This is a useful model as respiratory challenge induces both inflammatory cell recruitment and a robust T cell response^3, 10^. Furthermore, with 3-5 million severe cases yearly and an estimated 290,000-650,000 respiratory deaths according to WHO, IAV remains an important human pathogen.

Previous studies have demonstrated that innate immune cells such as natural killer cells and monocytes that are recruited to IAV-infected lungs express iCCRs during acute IAV viral infection^9, 11, 12^. Furthermore, CCR2 and CCR5 have been shown to play a role in recruitment and function of antigen-presenting cells (APCs) and T cells during IAV infection, and formation of the memory response^13-15^ thus linking these receptors to both innate and adaptive immunity. Here we have combined analysis of our two mouse strains to systematically explore the combinatorial roles of the iCCRs in the myeloid and T cell responses to IAV challenge. Overall, our data demonstrate that iCCRs are required for an optimal IAV-specific CD4 T cell response.

## RESULTS AND DISCUSSION

### Lung resident antigen presenting cells express combinations of iCCRs

We have previously shown that at rest, in the lung, iCCR KO mice display wild type levels of alveolar macrophages, dendritic cells and T cells indicating that these receptors are not involved in recruitment of these cells to the lung in homeostasis^8^. In contrast, inflammatory monocyte numbers were severely reduced. Notably, this reduction was substantially driven by CCR2 deficiency but the additional lack of CCR1, 3 and 5 in iCCR KO mice resulted in a further decrease of monocytes in naive lungs^8^. To look at this in more detail we have now used REP mice to document iCCR expression on APCs in the naive lung (Fig 1A, gating strategy Suppl Fig 1). Our data show that in line with previous findings^9^, >95% of monocytes express CCR2/mRuby2. The majority of recruited macrophages expressed CCR2, CCR5 or both. However, most alveolar macrophages and the majority of B cells did not express any iCCR, although a small population of each displayed CCR1 expression. Conventional DCs (cDCs) mainly displayed co-expression of CCR2 and CCR5. Thus iCCRs are abundantly expressed amongst antigen presenting cells in the naive lung.

**Figure 1:**
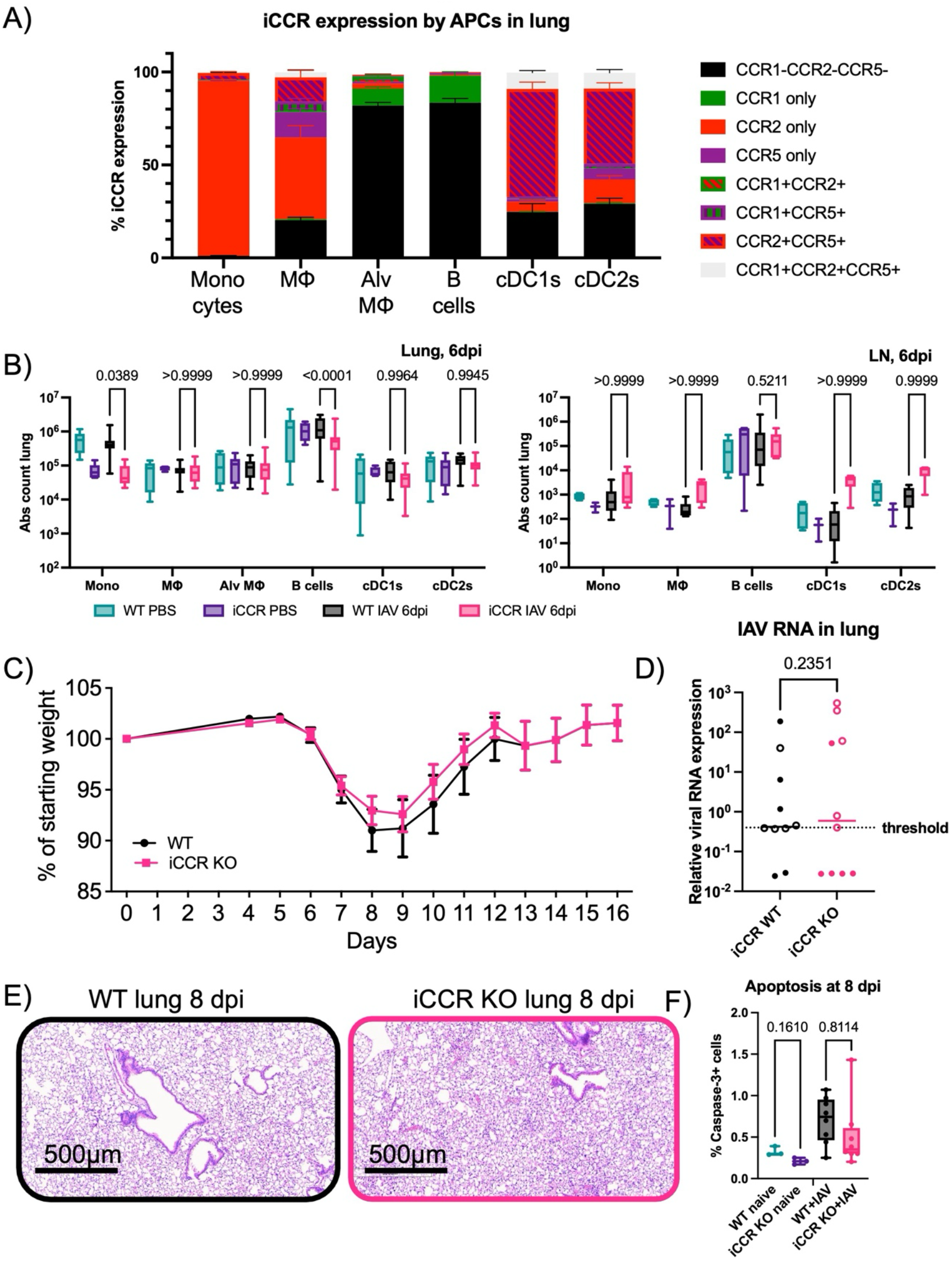
iCCRs have limited impact on IAV pathology and recruitment of APCs to the lung. REP, iCCR KO and matching WT mice were infected with 100-200 PFU WSN IAV or PBS. A) iCCR expression by antigen-presenting cells was determined in our REP mice using flow cytometry, matching gating strategy shown in Supp F1A-C. (n=5 REP mice and 1 matching WT for gating from 1 experiment, mean±SD) B) Absolute cell counts of antigen presenting cells at 6 days post infection in infected WT (black) or iCCR KO (pink) mice, compared to naive WT (teal) or iCCR KO (purple) mice (n=4-6 for naïve and 7-8 for infected mice, data pooled from two independent experiments, ‘min/max’ box plot). Gating strategy described in Supp Fig 1A-B. C) Collated weight curves for iCCR KO and WT mice (up to 7 dpi: WT n=17, iCCR KO n=30, 7-16 dpi WT n=9, iCCR KO n=17, collated data from 3 independent experiments, mean±SEM). D) Viral RNA expression as measured in lungs by qPCR at 8 dpi. (n=10/genotype, collated data from two independent experiments denoted by open and closed symbols, normalised to housekeeping with average WT IAV expression per experiment set to 1, line indicates median, two-sided T test. E) Histology at 8 dpi of WT (left, black) or iCCR KO (right, pink) mice (n=10 per genotype from two independent experiments, representative image shown). F) Quantified caspase 3 staining in lung at 8 dpi of naive and IAV-infected WT and iCCR KO mice (naive n=3, infected n=8 per genotype, example staining in Supp Fig 3).

### APC populations efficiently migrate to mLN and lung in the absence of iCCRs

Based on the above, we hypothesised that lack of iCCRs would result in a substantial decrease in migration of APCs to the draining lymph node after infection with IAV with a resultant reduction in the protective immune response. To investigate this, we quantified migration of key APCs in iCCR KO and WT mice infected with IAV strain WSN at 6 (data and gating in Suppl Fig 2) or 8 days post infection (dpi) in the lung and lung-draining mediastinal LN (mLN), (Fig 1B, gating strategy in Suppl Fig 1). We previously published a profound reduction of myelomonocytic cell recruitment in iCCR KO mice 8 days after IAV^8^. As expected, compared to wild-type mice, we indeed observed lower numbers of Ly6C^hi^ monocytes and CD64^+^ macrophages in the lungs of IAV-infected iCCR KO mice (Fig 1B, Supp Fig 1D and 2).

**Figure 2:**
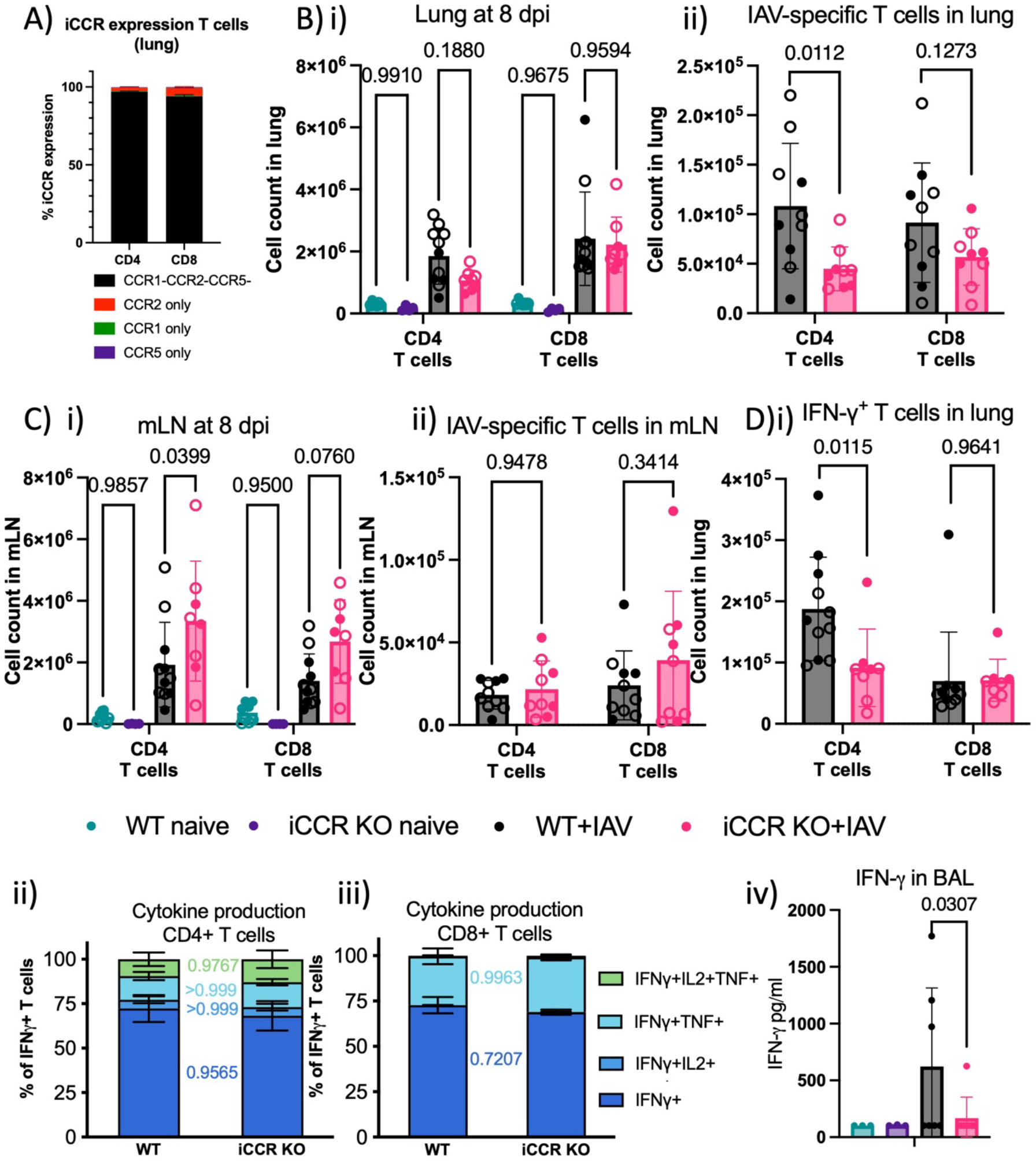
iCCRs play a role in development of IAV-specific CD4 T cells. REP, iCCR KO and WT mice were infected with 100-200 PFU WSN IAV, lung and mediastinal lymph node (mLN) were analysed at 8 days post infection. A) iCCR expression on lung T cells from uninfected REP mice quantified by flow cytometry (n=5 REP mice and 1 matching WT for gating, gating strategy in Supp Fig 4A-B) B-C) Absolute cell count of i) total and ii) IAV-specific CD4 and CD8 T cells in B) lung and C) mLN. Gates were set based on naive mice, as illustrated in Suppl Fig 4C. D) Cytokine production as measured by intracellular flow cytometry after challenge with peptide-loaded DCs, i) total number of IFN-γ^+^ CD4 and CD8 T cells and out of IFN-γ^+^ T cells percentages of multi-cytokine producing ii) CD4 and iii) CD8 T cells. Representative gating strategy in Suppl Fig 4D. Data visualized as mean±SD and analysed by T test with Welsh correction for 2 groups or 2-way ANOVA for >2 groups, n=8-11, data pooled from two independent experiments denoted by open and closed symbols.

After IAV infection, CD64^+^CD11c^+^ alveolar macrophages (AMs) were increased in WT compared to iCCR KO lungs This is in keeping with the literature as AMs are known to increase by two mechanisms; CCR1-mediated expansion of resident AMs^16^ and replenishment via CCR2-dependent recruitment from bone marrow-derived monocytes^17, 18^.

Interestingly, there was a slight decrease of B cells in the lung which may reflect the subset of B cells that express CCR1. We did not observe any differences in frequencies of cDC1s or cDC2s in the lung or mLN of wild type compared to iCCR KO mice in response to IAV infection (Fig 1B, Suppl Fig 2).

Others have reported that at earlier time points during more severe IAV infection models (up to 5 dpi after a high dose of X31), DC-specific CCR2-deficiency resulted in reduced recruitment of preDCs to the lung. Furthermore, CCR2 expression by these DCs was shown to be critical for formation of DC/T cell foci^13^. This difference in the context of our findings could relate to the timing, dose or strain of virus used. Indeed, we did not observe an increase in cell numbers of cDC1s or cDC2s, which is in line with previous work at 6 dpi after IAV infection with strain WSN^19^. It is curious though that whilst the majority of DCs do express CCR2 and CCR5, iCCR KO mice do not demonstrate a role for these receptors in the recruitment of DCs to the lung. Potentially CCR2 and CCR5 play a role in migration within the lung. This is in marked contrast to our previous study focusing on skin, demonstrating a central role for iCCRs in DC recruitment to the skin^8, 20^.

Overall, therefore, whilst APCs express iCCRs and a profound reduction in monocytes and macrophages is seen in iCCR KO mice, these receptors play no detectable role in controlling DC numbers in the lung or lung draining lymph node in this IAV model.

### iCCR deficiency has no impact on IAV-mediated pathology

Previous studies using mice deficient for single inflammatory chemokine receptors demonstrated that CCR2 and CCR5 have multiple roles during IAV infection that can exacerbate or ameliorate outcome, depending on IAV strain and dose^15, 21-23^. In the present study we did not observe any gross differences in IAV-mediated lung pathology in WT compared to iCCR KO mice. Specifically, there was no significant difference in weight loss between WT and iCCR KO mice, which is a measure of disease severity (Fig 1C). At 8 days post infection, which correlates to weight nadir and peak of the primary T cell response, we did not detect any difference between WT and iCCR KO mice in viral clearance as measured by viral RNA load (Fig 1D) or lung damage as visualised by histology (Fig 1E). Furthermore, we quantified histopathology by comparing caspase 3 staining, which visualises apoptotic cell death^24^, again demonstrating no difference between WT and iCCR KO mice (Fig 1F, Suppl Fig 3).

**Figure 3:**
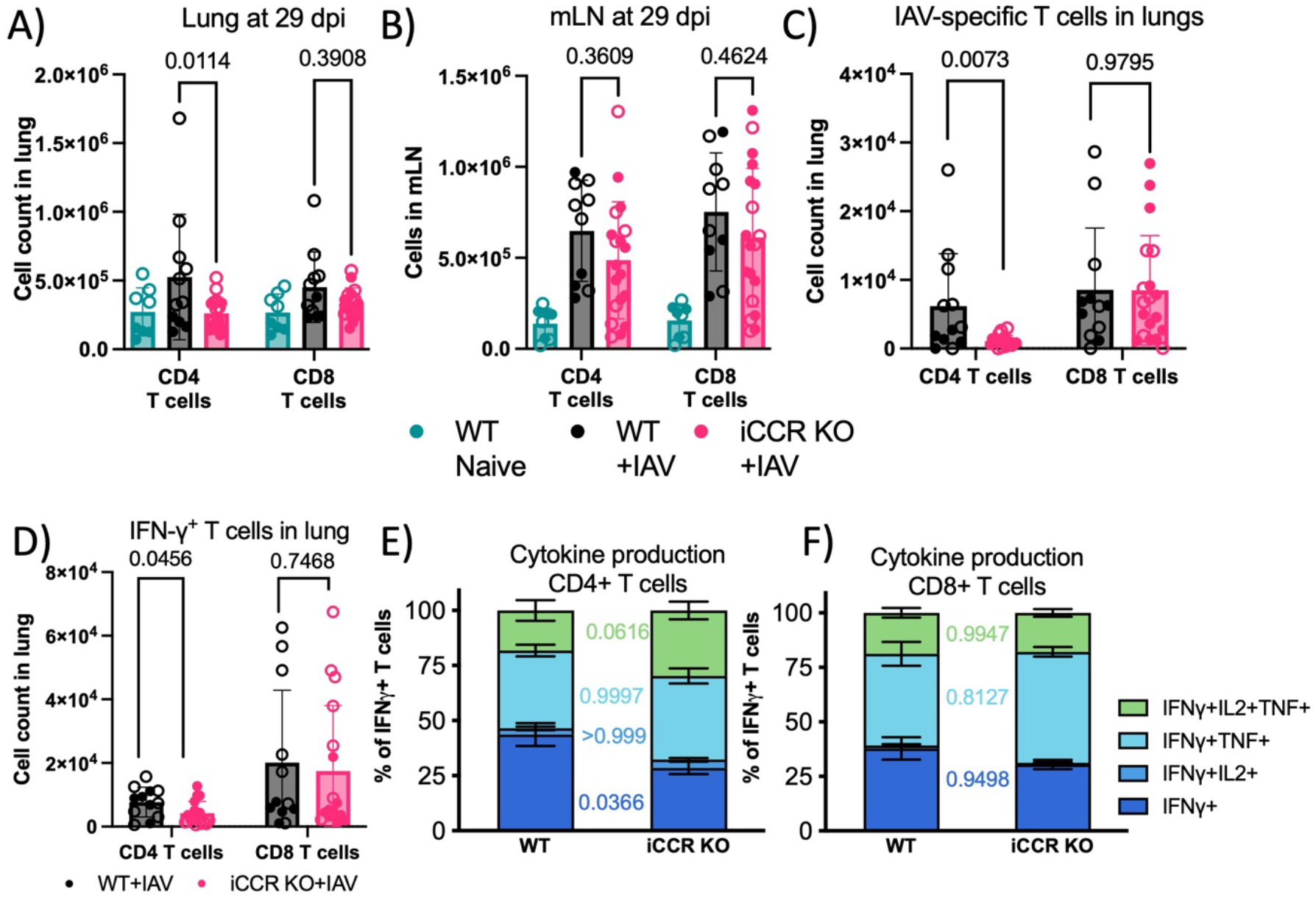
Defect in lung IAV-specific CD4 T cells remains during memory phase. iCCR KO and WT mice were infected with 100-200 PFU WSN IAV and at 29 days post infection T cells were analysed in lung and mediastinal lymph node (mLN) using flow cytometry. For gating strategy see Suppl Fig 4. CD4 and CD8 T cells were enumerated in A) lung and B) mLN. C) absolute cell count of IAV-specific T cells in lung as detected by IAV-specific tetramer staining. D-F) Cytokine production as measured by intracellular flow cytometry after challenge with peptide-loaded DCs in CD4 and CD8 T cells. D) total CD4 and CD8 IFN-γ^+^ T cells in lung, with IFN-γ^+^ cells further analysed for E-F) TNF-α and IL-2 production. Data visualized as mean±SD, and analysed by T test with Welsh correction for 2 groups or 2-way ANOVA for >2 groups, n=8-17 with data pooled from two independent experiments denoted by open and closed symbols.

Overall, these results demonstrate that despite extensive expression of iCCRs in leucocytes of importance in the immune/inflammatory response to IAV challenge, they have no detectable combined role in the lung pathology arising in response to IAV in our model.

### iCCRs play a significant role in the generation of IAV-specific CD4 T cells

We next investigated the role of iCCRs in T cell migration and activation in response to IAV challenge. Using REP mice, we initially examined iCCR expression by lung T cells. In line with our previous findings using inflammatory models^9^, for naive mice only a small minority of lung T cells expressed CCR2 with no evidence of expression of any other iCCRs by this cell type (Fig 2A, gating in Suppl Fig 4A-B).

Comparing naive WT and iCCR KO mice, at 8 dpi we observed substantial recruitment of CD4 and CD8 T cells to the lung which was similar in both WT and iCCR KO mice (Fig 2Bi). However, the number of IAV-specific CD4, but not CD8, T cells as measured by IAV-tetramer-binding was significantly decreased in lungs of iCCR KO mice (Figure 2Bii, gating in Suppl Fig 4C).

We detected significantly more total CD4 T cells in the mLN at 8 dpi (Figure 2Ci), whilst we did not see a significant accumulation of IAV-specific CD4 T cells in the mLN (Fig 2Cii).

The similar number of IAV-specific CD4 T cells in the mLN demonstrates that T cell activation has not been affected by the absence of iCCR. In contrast, the reduction in the numbers of IAV-specific CD4 T cells in the lungs of iCCR KO mice suggests that iCCRs are involved in IAV-specific CD4 T cell recruitment. Alternative explanations for this difference include altered differentiation into lung-homing CD4 T cells and/or reduced survival of the cells once recruited to the lung.

A key effector role for T cells is mediated through the production of cytokines such as IFN-γ which stimulates an antiviral immune response^25^. IFN-γ is an antiviral cytokine that links the innate and adaptive immune response via several mechanisms, including upregulation of major histocompatibility class I (MHC-I)^26^ and MHC-II^27^ for antigen presentation and upregulation of many chemokines including multiple iCCR ligands for immune cell recruitment^25^. NK cells and T cells are the main producers of IFN-γ during IAV infection^28^. Production of IFN-γ by T cells is therefore a marker of immune activation. As such we next investigated IFN-γ production by T cells using intracellular staining following *in vitro* culture with IAV nucleoprotein (IAV-NP) immunodominant epitopes (gating in Suppl Fig 4D). In line with the decreased number of IAV-specific CD4 T cells in the lung, we observed lower numbers of IFN-γ-producing CD4 but not CD8 T cells (Fig 2Di). When we further characterised these IFN-γ^+^ T cells we did not detect any difference in expression of two other key cytokines implicated in protection against IAV, TNF-α and IL-2^29, 30^ in WT compared to iCCR KO mice (Fig 2Dii-iii). This reduced IFN-γ production by CD4 T cells was reflected in a reduced level of IFN-γ in bronchial alveolar lavage (BAL) as measured by Luminex (Fig 2Div).

Thus, these data demonstrate an important role for iCCRs in the CD4 T cell response to IAV infection.

### The defect in IAV-specific CD4 T cells is maintained during memory phase

To determine whether the observed defect in IAV-specific and IFN-γ-producing CD4 T cells is maintained over time, we infected WT and iCCR KO mice and quantified APC subsets at 29 dpi (Suppl Fig 5A, gating in Suppl Fig 2). At this stage, alveolar and monocyte-derived macrophage numbers were similar to matching, uninfected WT mice. There were no differences between WT and iCCR KO mice in the various quantified DC subsets in lung or mLN. However, in the absence of iCCRs, the number of B cells was slightly albeit statistically significantly lower in the lung whilst higher in the mLN at this time point which may indicate a minor contribution of CCR1 to overall B cell activity in response to IAV infection.

After the acute IAV infection is cleared, T cell populations contract and memory T cells remain present in the lung and mLN^28, 31^. Indeed, the total number of T cells in the lung and mLN at 29 dpi had decreased by 3-4 fold compared to 8 dpi but in mLN remained higher than naive WT controls (Suppl Fig 5C). Again, we observed that CD8 T cells were unaffected by iCCR deficiency, whilst CD4 T cells remained significantly decreased in the lungs of iCCR KO mice but not mLN at 29 dpi (Fig 3A-B). In line with the numbers of IAV-specific T cells during acute infection, in the absence of iCCRs the number of IAV-specific CD4 T cells in the lung as measured by IAV-specific tetramer staining remained substantially reduced at 29 dpi, whilst CD8 T cells were similar between WT and iCCR KO animals (Fig 3C).

Subsequently, we determined the capacity of T cells to produce inflammatory cytokines at 29 dpi in response to IAV NP epitopes (Fig 3D-F). The absolute number of IFN-γ^+^ CD4 T cells remained substantially decreased in the lungs of iCCR KO mice, whilst numbers of IFN-γ^+^ CD8 T cells were similar in lungs of WT and iCCR KO mice at this memory time point (Fig 3D). In line with our findings during acute infection, there was no difference in multi-functional cytokine production by CD8 T cells in the lung. In contrast, out of IFN-γ^+^ CD4 T cells a lower percentage of single IFN-γ+ cells were present in iCCR KOs as compared to the WT mice, whilst there was a trend towards increased IFN-γ^+^/IL-2/TNF-α positive CD4 T cells in the absence of iCCR (Fig 3E-F). A possible explanation for this could be a difference in survival and proliferation by the different subsets of CD4 T cells; single-cytokine producing CD4 T cells are more proliferative but less likely to survive long-term compared to the triple-cytokine producing CD4 T cells^31^. Therefore, the slightly increased proportion of triple-cytokine producing CD4 T cells in iCCR KO mice may have compensated for the decrease in IAV-specific CD4 T cells. This could be driven by the importance of CCR5 in the nanoclustering of T cell receptors on memory CD4 T cells^32, 33^.

To summarise, despite extensive expression of iCCRs on APCs and a marked reduction of alveolar macrophages, recruited monocytes and macrophages we observed no impact of iCCR on viral pathology during a primary IAV model. These changes did not impact the IAV-specfic CD8 T cell response nor on the activation of IAV-specific CD4 T cells in the mLN. We did find a reduced IAV-specific lung CD4 T cell response in the absence of iCCR. This reduction at both primary and memory time points following IAV infection could have been due to a knock-on effect of the altered innate response and/or reduced migration to, or survival within, the lung of IAV-specific CD4 T cells.

## ACKNOWLEGEMENTS

We would like to acknowledge Georgia Perona-Wright and Ed Roberts for stimulating discussion; Alistair Gamble, Carmen San Martin Martinez, Pietro Cocchiara, Jack Jones, Patrick Shearer and Emma Nilsson for technical assistance; the University of Glasgow Shared Research Facilities Cellular Analysis and Biological Services for their support and assistance.

This study was funded by a Wellcome Trust Investigator Award (217093/Z/19/Z) and an MRC Programme Grant (MR/V010972/1).

---

**Supplementary Figure 1: APC gating strategy for naive and six days post infection data in Figure 1**.

A) Gating strategy used for APCs in lung at 6 days post infection in Fig 1A-B.

B) Table summarising markers expressed by the various described subsets of cells in the lung, based on gating strategy in A) and used for Figure 1A-B.

C) Gating strategy for reporter proteins, with WT control in black and representative REP sample in pink. Cells were gated as in A), with cell subsets gated for CCR1/CCR2 and every subset subsequently gated for CCR5 expression. In line with our previous publication, no CCR3 was observed on the analysed cell types^9^.

D) Representative tSNE plots of flow cytometric characterization of antigen presenting cells with quantification as percentage of live cells in the lung at 6 dpi (Pooled data from 2 independent experiments with 4-8 mice per group, error bars±SEM).

**Supplementary Figure 2: Gating and APC quantification at eight days post infection**.

A) Gating strategy used for APCs in lung at 8 days post infection in B).

B) Absolute cell counts of antigen presenting cells at 8 days post infection in infected WT (black) or iCCR KO (pink) mice, compared to naive WT (teal) mice (n=8-11, data pooled from two independent experiments, ‘min/max’ box plot).

**Supplementary Figure 3: Representative images for caspase 3 staining at eight days post infection**.

Example images from caspase 3 stain from lungs of naive and IAV-infected WT and iCCR KO mice at 8 dpi matching quantification in Fig 1F. Arrows indicate positive cells.

**Supplementary Figure 4: T cell and reporter gating strategy**.

A) Gating strategy for CD4 and CD8 T cells with representative images from lung gating used in Figure 2A.

B) Gating strategy for iCCR reporter mice in Figure 2A. Overlay of T cells from WT in black and REP in colour.

C) Gating strategy for IAV NP tetramer staining (Fig 2 and 3). Cells were pregated for live, non-doublet, TCR-β^+^, DUMP^-^ (B220/MHC-II/F4/80) NK1.1^-^ CD4 or CD8 T cells before final gate of CD44^+^ NP IAV^+^. Gates for NP-positive cells were set using naïve mice as controls.

D) T cells were pregated for live, non-doublet, DUMP^-^, CD4 and CD8 T cells as in C) before gating for CD44^+^ IFN-γ^+^ cells, which were further subsetted for TNF-α and IL-2 production.

**Supplementary Figure 5: Quantification of T cells and APCs in lung and mLN**.

A) Absolute cell counts of antigen presenting cells at 29 (B) days post infection in infected WT (black) or iCCR KO (pink) mice, compared to naive WT (teal) mice in lung and mLN.

B) Total number of T cells defined as TCR-αβ-positive cells in lung and lymph node, and C) IAV-specific CD4 and CD8 T cells as proportion of total T cells at 8 days post infection in infected WT (black) or iCCR KO (pink) mice, compared to naive WT (teal) and iCCR KO mice (purple).

D) Total number of T cells defined as TCR-αβ-positive cells in lung and lymph node, and E) IAV-specific CD4 and CD8 T cells as proportion of total T cells at 29 days post infection.

Data visualized as mean±SD, and analysed by T test with Welsh correction for 2 groups or 2-way ANOVA for >2 groups, n=8-17 with data pooled from two independent experiments denoted by open and closed symbols.

## METHODS

### Animals

Transgenic mouse strains REP^9^, iCCR KO and matching wild types (both on C57BL/6 background)^8^ were bred in house in individually ventilated cages in a specific pathogen-free facility. For all experiments 10-week-old males were used. Mice were housed per genotype but where possible researchers were blinded to infection status until analysis was completed. Sample sizes were based on previous studies. When required to move mice, they were acclimatised for 2-7 days. All breeding and procedures were approved by local ethics and the UK home office (license PP6655603).

### Influenza infection

IAV/WSN/33 (H1N1) was prepared and titrated in MDCK cells per standard protocols^34^. Mice were briefly anesthetised with isoflurane before inoculation of 20 ul PBS or virus diluted in PBS. Virus inoculum was adapted to mouse weight at infection day; 100 plaque forming units (PFU) for mice <20 gram, 150 PFU for 20-25 gram and 200 PFU for >25 gram body weight.

Infected mice were excluded from the study if they lost >20% of their starting body weight which was predefined as a humane endpoint or if the lung-draining mediastinal lymph node was not enlarged at time of cull.

For 8 and 29 dpi, IAV-infected WT and iCCR KO mice were compared to each other and naive WT mice. For 6 dpi, WT and iCCR KO mice were either IAV or mock infected. To prevent confounding factors, treatment was predetermined randomly using Microsoft Excel and groups were infected in random order.

### *Ex vivo* reactivation

Single cell suspensions from lungs containing T cells were reactivated by co-culturing with murine bone marrow-derived GM-CSF DCs that were exposed to IAV antigen or peptide as previously described^34^ at an approximate ratio of 10 T cells to 1 DC in the presence of GolgiPlug (BD Biosience) for six hours at 37°C 5% CO_2_. Cells were subsequently stained for flow cytometry with B220 (RA3-6B2) and MHC-II (M5114, both fitc for dump), CD4-APC-e780 (RMRM4-5), CD8-PeCy7 (53-6.7) and CD44-PerCP-Cy5.5 (IM7) before intracellular staining using cytofix/cytoperm (BD Bioscience) as per manufacturer’s instructions and IL-2-APC (JES6-5H4), TNF-α-FITC (MP6-XT22) and IFN-γ-Pe (XMG1.2).s

**Flow cytometry** At 6, 8 or 29 days post infection, mice were culled, perfused with PBS and tissues harvested. Mediastinal lymph node and lungs were mechanically disrupted and digested in 1.3 mg collagenase D (Roche) and 30 ug/ml DNase (Sigma) for 40 minutes at 37°C in a shaking incubator. Red blood cells in lung samples were lysed using ACK lysis solution (ThermoFisher).

Single cell suspensions were stained for viability using fixable viability stain e506 (eBioscience) and blocked using FcR blocking reagent (Miltenyi) before staining with B220-Fitc (RA3-6B2), TCR-PerCP-Cy5.5 (H57-597), CD11c-APC-Cy7 (N418), CD64-BV421 (X54-5/7.1), SiglecH-Pe (551), CD11b-PeCy7 (M1/70), MHC-II-AlexaFluor700 (M5/114.15.2), CD103-APC (M290) (all Biolegend).

Alternatively, cells were stained with PE or APC-labelled tetramers (NIH tetramer core, IA^b^/NP_311-325_ or D^b^/NP_368-374_) for 2 hours at 37°C in RPMI/10% fetal calf serum/100ug/ml penicillin-streptomycin/2 mM L-glutamine/Fc block, with the following antibody cocktail added for the final 30 minutes: NK1.1-BV421 (PK136), CD44-APC-e780 (IM7), CD8-PeCy7 (53-6.7), B220 (RA3-6B2)/MHC-II (M5/114.15.2)/F4/80 (BM8)-Fitc (dump), TCR-PerCP-Cy5.5 (H57-597) and CD4-AF700 (RM4-5) before staining with fixable viability stain e506.

Cells were acquired on a BD FACSCelesta, BD LSR or BD Fortessa and analysed using FlowJo v10. Where data is collated, the same analyser was used to obtain data.

### Histology

At 8 dpi, mice were perfused with 10 mL PBS containing 5 mM EDTA through the heart. Lungs were then inflated from the trachea using 10% formalin, tied off using thread before lungs were dissected out and placed in container with 10% formalin at 4°C. After overnight incubation, 10% formalin was replaced by 70% Ethanol and blinded samples were processed in paraffin and Haematoxylin and Eosin staining or caspase-3 staining by University of Glasgow Histology Research Service. Whole slides were scanned at 40x magnification.

For quantification, a blinded researcher selected five random field of views per section, and counted the number of caspase-3 positive cells and the total cell number to calculate the percentage of caspase-3 positive cells.

### qPCR

After perfusion, tissues were fixed in RNAlater overnight at 4°C and kept at - 70°C until RNA extraction using TRIzol and RNA purification kit (Invitrogen), with additional on column DNase step (PureLink DNase, Invitrogen). BAL was directly frozen at -70°C until RNA extraction with QIAamp Viral RNA mini kit (Qiagen) according to manufacturer’s protocol.

RNA was converted to cDNA using High-Capacity cDNA Reverse Transcription kit and analysed by qPCR using Perfecta SYBR, all procedures as described previously^35^. Primer sequences: Standard primers for housekeeping gene m18S 5’-CGTAGTTCCGACCATAAACGA-3’ and 5’-ACATCTAAGGGCATCACAGACC-3’ and Influenza NP 5’-TGCACCGAACTCAAACTCAG-3’ and 5’-TTGCATCAGAGAGCACATCC-3’, qPCR primers for m18S 5’-GACTCAACACGGGAAACCTC-3’ and 5’-TAACCAGACAAATCGCTCCAC-3’, for influenza NP 5’-CTCGTCGCTTATGACAAAGAAG -3’ and 5’-AGATCATCATGTGAGTCAGAC-3’.

### Luminex

Cytokine expression in BAL was analysed using a customised Magnetic Luminex Multiplex assay (R&D Systems) and analysed on a Luminex 200 machine (Biorad).

### Analysis

Data were analysed in Prism10 using one or two-way ANOVA with multiple comparisons or Welch’s T tests as described in the figure legends. Single animals were used as the experimental unit. Open and closed circles in dotplots indicate animals from independent experiments.

## REFERENCES

1. Holt PG, Strickland DH, Wikstrom ME, Jahnsen FL. Regulation of immunological homeostasis in the respiratory tract. Nat Rev Immunol 2008; 8(2): 142–152.

2. Ardain A, Marakalala MJ, Leslie A. Tissue-resident innate immunity in the lung. Immunology 2019; 159(3): 245–256.

3. Alon R, Sportiello M, Kozlovski S, Kumar A, Reilly EC, Zarbock A et al. Leukocyte trafficking to the lungs and beyond: lessons from influenza for COVID-19. Nature Reviews Immunology 2020; 21(1): 49–64.

4. Bachelerie F, Ben-Baruch A, Burkhardt AM, Combadiere C, Farber JM, Graham GJ et al. International Union of Basic and Clinical Pharmacology. LXXXIX. Update on the Extended Family of Chemokine Receptors and Introducing a New Nomenclature for Atypical Chemokine Receptors. Pharmacological Reviews 2014; 66(1): 1–79.

5. Griffith JW, Sokol CL, Luster AD. Chemokines and chemokine receptors: positioning cells for host defense and immunity. Annu Rev Immunol 2014; 32: 659–702.

6. Rot A, von Andrian UH. Chemokines in innate and adaptive host defense: basic chemokinese grammar for immune cells. Annu Rev Immunol 2004; 22: 891–928.

7. Nomiyama H, Osada N, Yoshie O. A family tree of vertebrate chemokine receptors for a unified nomenclature. Dev Comp Immunol 2011; 35(7): 705–715.

8. Dyer DP, Medina-Ruiz L, Bartolini R, Schuette F, Hughes CE, Pallas K et al. Chemokine Receptor Redundancy and Specificity Are Context Dependent. Immunity 2019; 50(2): 378-389.e375.

9. Medina-Ruiz L, Bartolini R, Wilson GJ, Dyer DP, Vidler F, Hughes CE et al. Analysis of combinatorial chemokine receptor expression dynamics using multi-receptor reporter mice. Elife 2022; 11.

10. Bender BS, Croghan T, Zhang L, Small PA. Transgenic mice lacking class I major histocompatibility complex-restricted T cells have delayed viral clearance and increased mortality after influenza virus challenge. The Journal of experimental medicine 1992; 175(4): 1143–1145.

11. Ferrero MR, Tavares LP, Garcia CC. The Dual Role of CCR5 in the Course of Influenza Infection: Exploring Treatment Opportunities. Frontiers in Immunology 2022; 12.

12. Brownlie D, Rødahl I, Varnaite R, Asgeirsson H, Glans H, Falck-Jones S et al. Comparison of Lung-Homing Receptor Expression and Activation Profiles on NK Cell and T Cell Subsets in COVID-19 and Influenza. Frontiers in Immunology 2022; 13.

13. Cabeza-Cabrerizo M, Minutti CM, da Costa MP, Cardoso A, Jenkins RP, Kulikauskaite J et al. Recruitment of dendritic cell progenitors to foci of influenza A virus infection sustains immunity. Sci Immunol 2021; 6(65): eabi9331.

14. Kohlmeier JE, Miller SC, Smith J, Lu B, Gerard C, Cookenham T et al. The Chemokine Receptor CCR5 Plays a Key Role in the Early Memory CD8+ T Cell Response to Respiratory Virus Infections. Immunity 2008; 29(1): 101–113.

15. Lin KL, Suzuki Y, Nakano H, Ramsburg E, Gunn MD. CCR2+ Monocyte-Derived Dendritic Cells and Exudate Macrophages Produce Influenza-Induced Pulmonary Immune Pathology and Mortality. The Journal of Immunology 2008; 180(4): 2562–2572.

16. Tapmeier TT, Howell JH, Zhao L, Papiez BW, Schnabel JA, Muschel RJ et al. Evolving polarisation of infiltrating and alveolar macrophages in the lung during metastatic progression of melanoma suggests CCR1 as a therapeutic target. Oncogene 2022; 41(46): 5032–5045.

17. Li F, Piattini F, Pohlmeier L, Feng Q, Rehrauer H, Kopf M. Monocyte-derived alveolar macrophages autonomously determine severe outcome of respiratory viral infection. Sci Immunol 2022; 7(73): eabj5761.

18. Mould KJ, Barthel L, Mohning MP, Thomas SM, McCubbrey AL, Danhorn T et al. Cell Origin Dictates Programming of Resident versus Recruited Macrophages during Acute Lung Injury. American Journal of Respiratory Cell and Molecular Biology 2017; 57(3): 294–306.

19. Hargrave KE, Worrell JC, Pirillo C, Brennan E, Masdefiol Garriga A, Gray JI et al. Lung influenza virus-specific memory CD4 T cell location and optimal cytokine production are dependent on interactions with lung antigen-presenting cells. Mucosal Immunology 2024.

20. Bartolini R, Medina-Ruiz L, Hayes AJ, Kelly CJ, Halawa HA, Graham GJ. Inflammatory Chemokine Receptors Support Inflammatory Macrophage and Dendritic Cell Maturation. ImmunoHorizons 2022; 6(11): 743–759.

21. Dawson TC, Beck MA, Kuziel WA, Henderson F, Maeda N. Contrasting Effects of CCR5 and CCR2 Deficiency in the Pulmonary Inflammatory Response to Influenza A Virus. The American Journal of Pathology 2000; 156(6): 1951–1959.

22. Lin S-J, Lo M, Kuo R-L, Shih S-R, Ojcius DM, Lu J et al. The pathological effects of CCR2+ inflammatory monocytes are amplified by an IFNAR1-triggered chemokine feedback loop in highly pathogenic influenza infection. Journal of Biomedical Science 2014; 21(1).

23. Carlin LE, Hemann EA, Zacharias ZR, Heusel JW, Legge KL. Natural Killer Cell Recruitment to the Lung During Influenza A Virus Infection Is Dependent on CXCR3, CCR5, and Virus Exposure Dose. Frontiers in Immunology 2018; 9.

24. Wurzer WJ, Planz O, Ehrhardt C, Giner M, Silberzahn T, Pleschka S et al. Caspase 3 activation is essential for efficient influenza virus propagation. EMBO J 2003; 22(11): 2717–2728.

25. Rauch I, Muller M, Decker T. The regulation of inflammation by interferons and their STATs. JAKSTAT 2013; 2(1): e23820.

26. Shirayoshi Y, Burke PA, Appella E, Ozato K. Interferon-induced transcription of a major histocompatibility class I gene accompanies binding of inducible nuclear factors to the interferon consensus sequence. Proc Natl Acad Sci U S A 1988; 85(16): 5884–5888.

27. Steimle V, Siegrist CA, Mottet A, Lisowska-Grospierre B, Mach B. Regulation of MHC class II expression by interferon-gamma mediated by the transactivator gene CIITA. Science 1994; 265(5168): 106–109.

28. Finney GE, Hargrave KE, Pingen M, Purnell T, Todd D, MacDonald F et al. Triphasic production of IFNγ by innate and adaptive lymphocytes following influenza A virus infection. Discovery Immunology 2023; 2(1).

29. Alam F, Singh A, Flores-Malavet V, Sell S, Cooper AM, Swain SL et al. CD25-Targeted IL-2 Signals Promote Improved Outcomes of Influenza Infection and Boost Memory CD4 T Cell Formation. J Immunol 2020; 204(12): 3307–3314.

30. DeBerge MP, Ely KH, Enelow RI. Soluble, but Not Transmembrane, TNF-α Is Required during Influenza Infection To Limit the Magnitude of Immune Responses and the Extent of Immunopathology. The Journal of Immunology 2014; 192(12): 5839–5851.

31. Westerhof LM, Noonan J, Hargrave KE, Chimbayo ET, Cheng Z, Purnell T et al. Multifunctional cytokine production marks influenza A virus-specific CD4 T cells with high expression of survival molecules. Eur J Immunol 2023: e2350559.

32. Martín-Leal A, Blanco R, Casas J, Sáez ME, Rodríguez-Bovolenta E, de Rojas I et al. CCR 5 deficiency impairs CD4+ T-cell memory responses and antigenic sensitivity through increased ceramide synthesis. The EMBO Journal 2020; 39(15).

33. Blanco R, Gomez de Cedron M, Gamez-Reche L, Martin-Leal A, Gonzalez-Martin A, Lacalle RA et al. The Chemokine Receptor CCR5 Links Memory CD4(+) T Cell Metabolism to T Cell Antigen Receptor Nanoclustering. Front Immunol 2021; 12: 722320.

34. Westerhof LM, McGuire K, MacLellan L, Flynn A, Gray JI, Thomas M et al. Multifunctional cytokine production reveals functional superiority of memory CD4 T cells. Eur J Immunol 2019; 49(11): 2019–2029.

35. Lefteri DA, Bryden SR, Pingen M, Terry S, McCafferty A, Beswick EF et al. Mosquito saliva enhances virus infection through sialokinin-dependent vascular leakage. Proc Natl Acad Sci U S A 2022; 119(24): e2114309119.

