## Supplemental Figures for "Inflammatory chemokine receptors CCR1, CCR2, CCR3 and CCR5 are essential for an optimal T cell response to influenza"

**Supplementary Figure 1. APC gating strategy for naive and six days post infection data in Figure 1**

A)

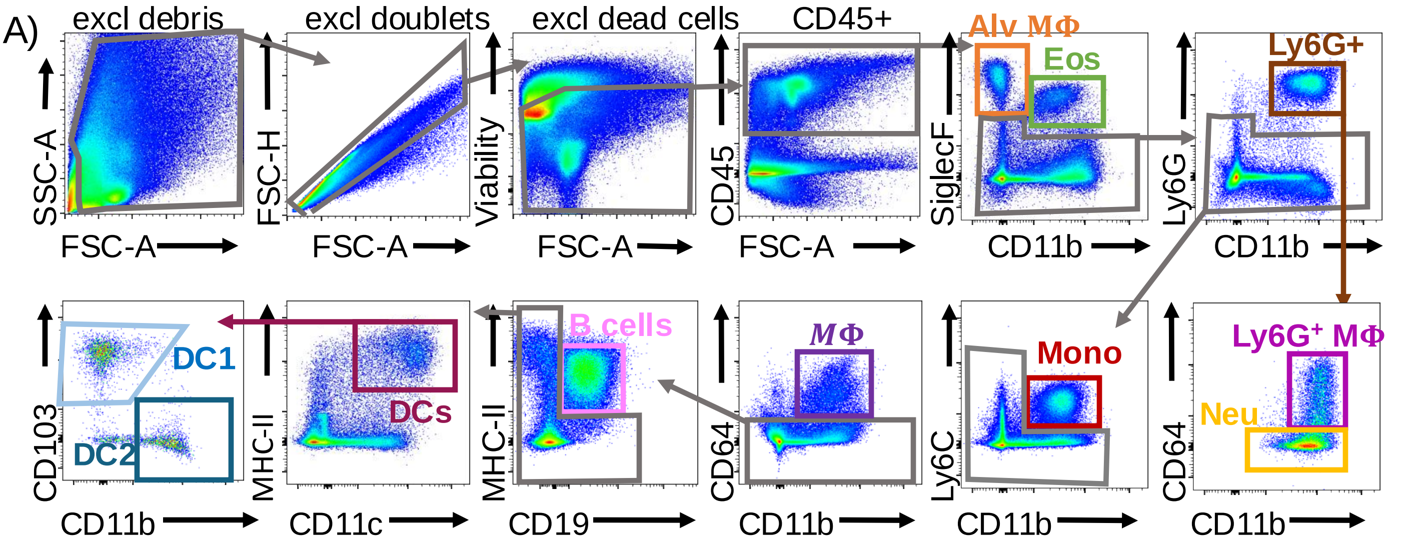


B)

| Subset | SiglecF | CD11b | Ly6G | CD64 | Ly6C | MHC-II | CD19 | CD11c | CD103 |
|---|---|---|---|---|---|---|---|---|---|
| Alveolar macrophage | high | - | | | | | | | |
| Eosinophil | + | mid | | | | | | | |
| Ly6G$^{+}$ macrophage | - | + | + | + | | | | | |
| Neutrophil | - | + | + | - | | | | | |
| Monocyte | - | + | - | | + | | | | |
| Macrophage | - | + | - | + | - | | | | |
| B cells | - | | - | - | - | + | + | | |
| DC1 | - | | - | - | - | high | | + | + |
| DC2 | - | + | - | - | - | high | | + | |

C)

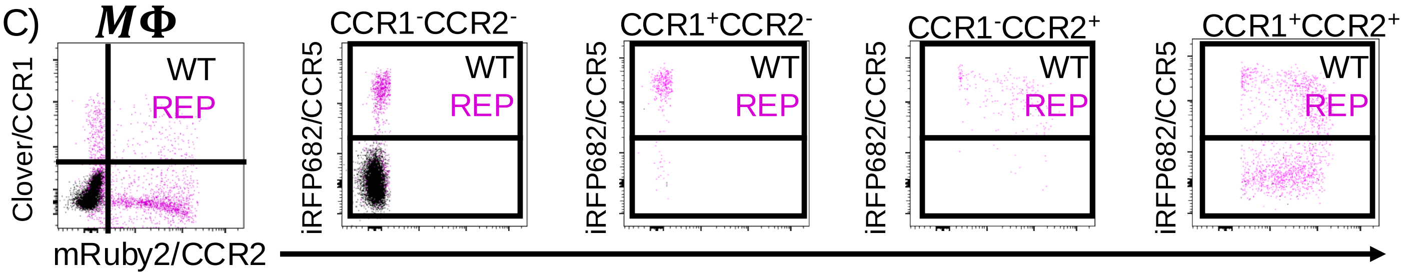


D)

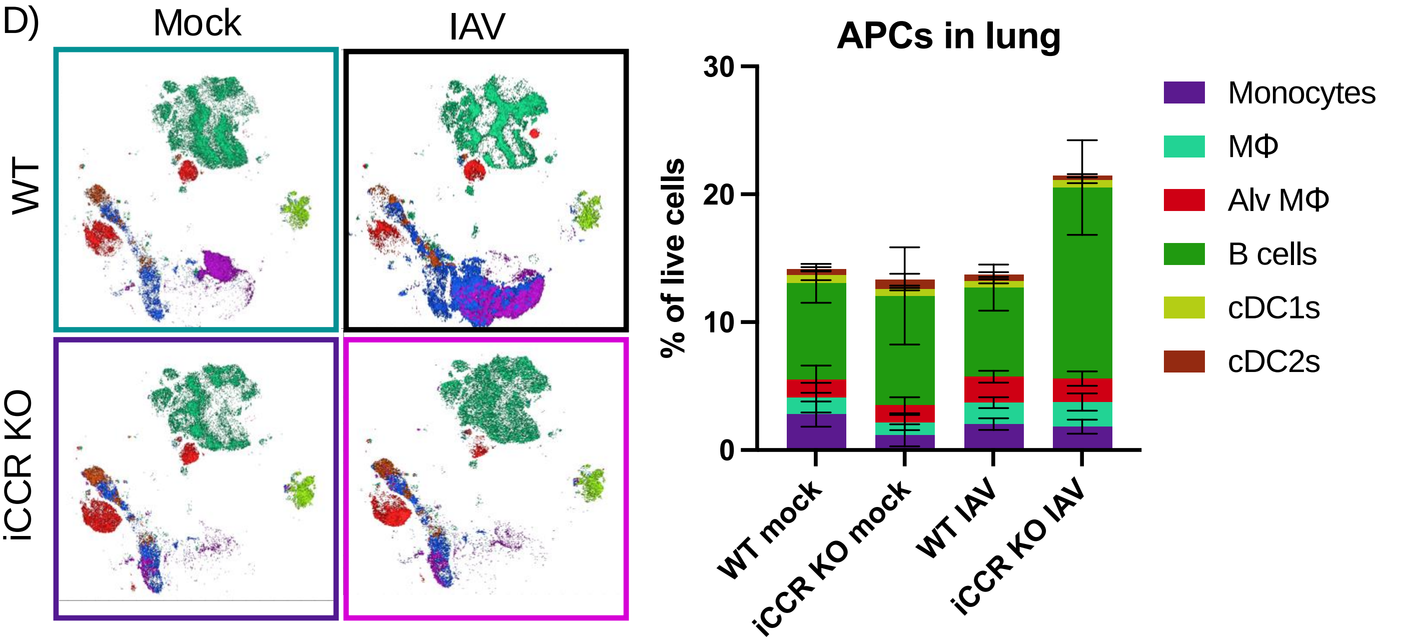

### Supplementary Figure 2: Gating and APC quantification at eight days post infection.

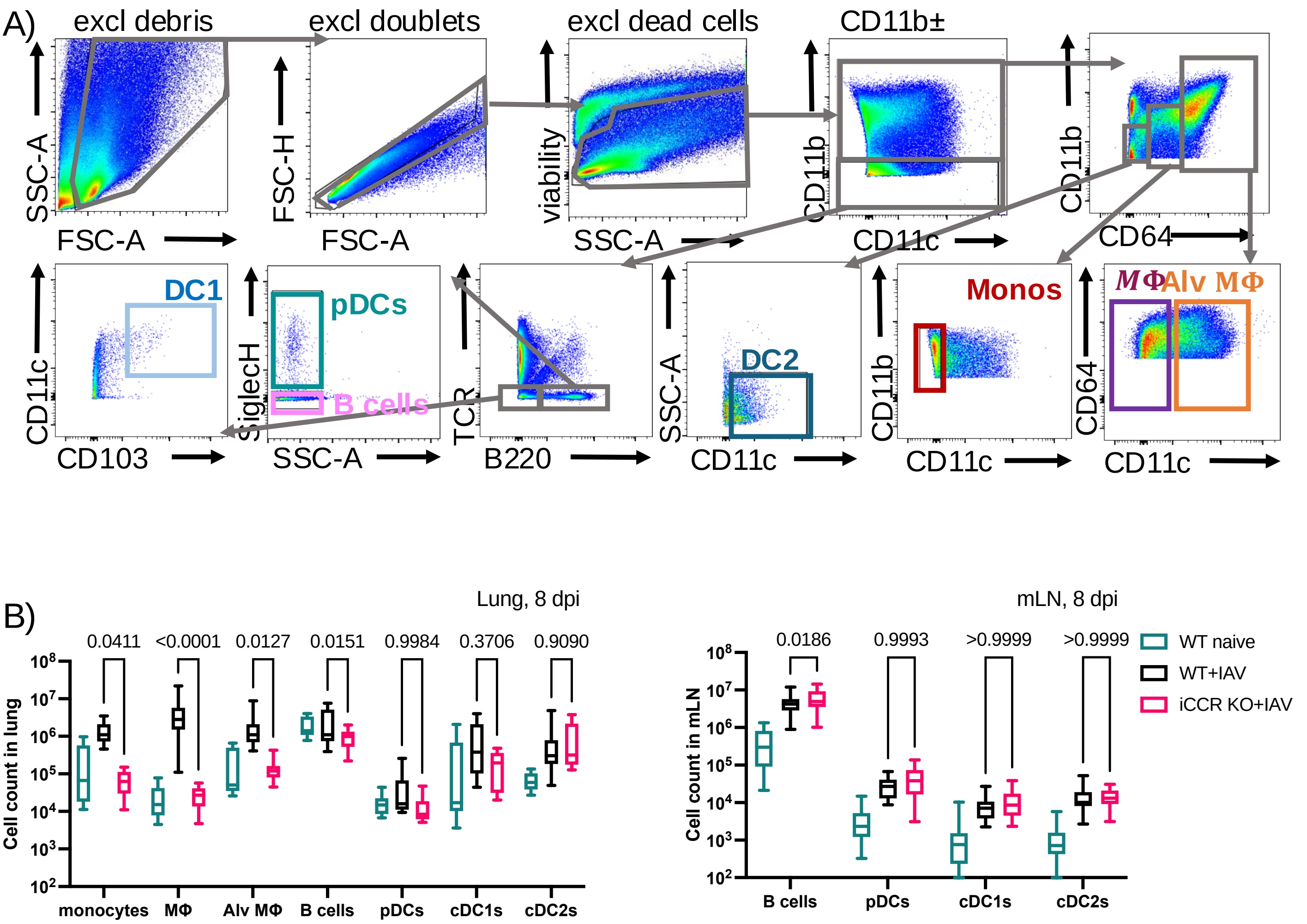

**Supplementary Figure 3: Representative images for caspase 3 staining at eight days post infection.**

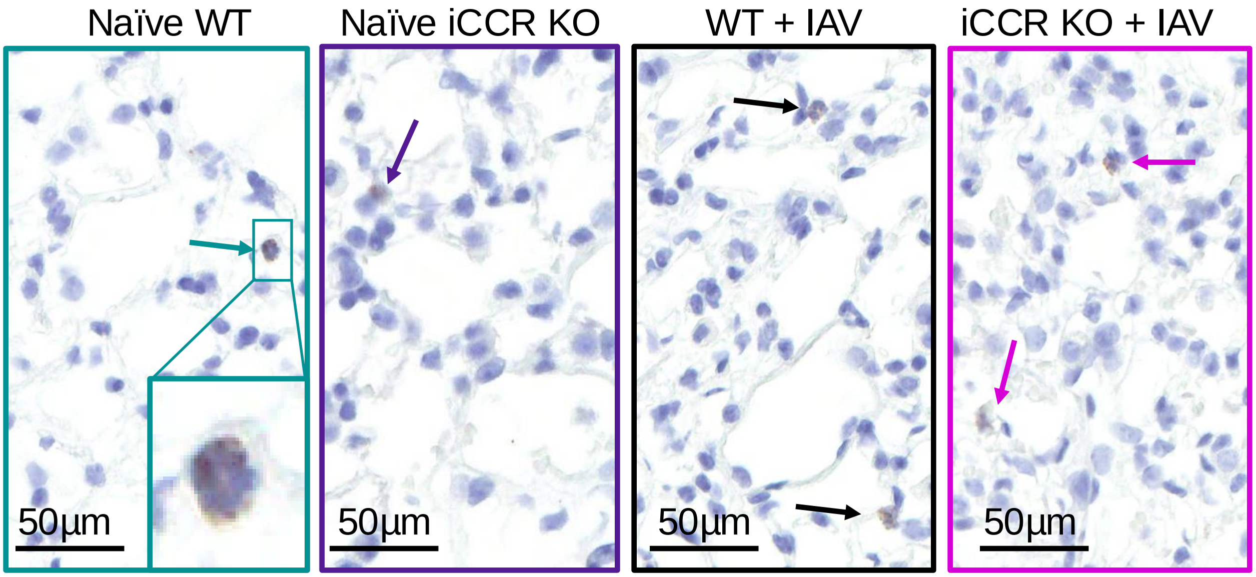

### Supplementary Figure 4. T cell and reporter gating strategy

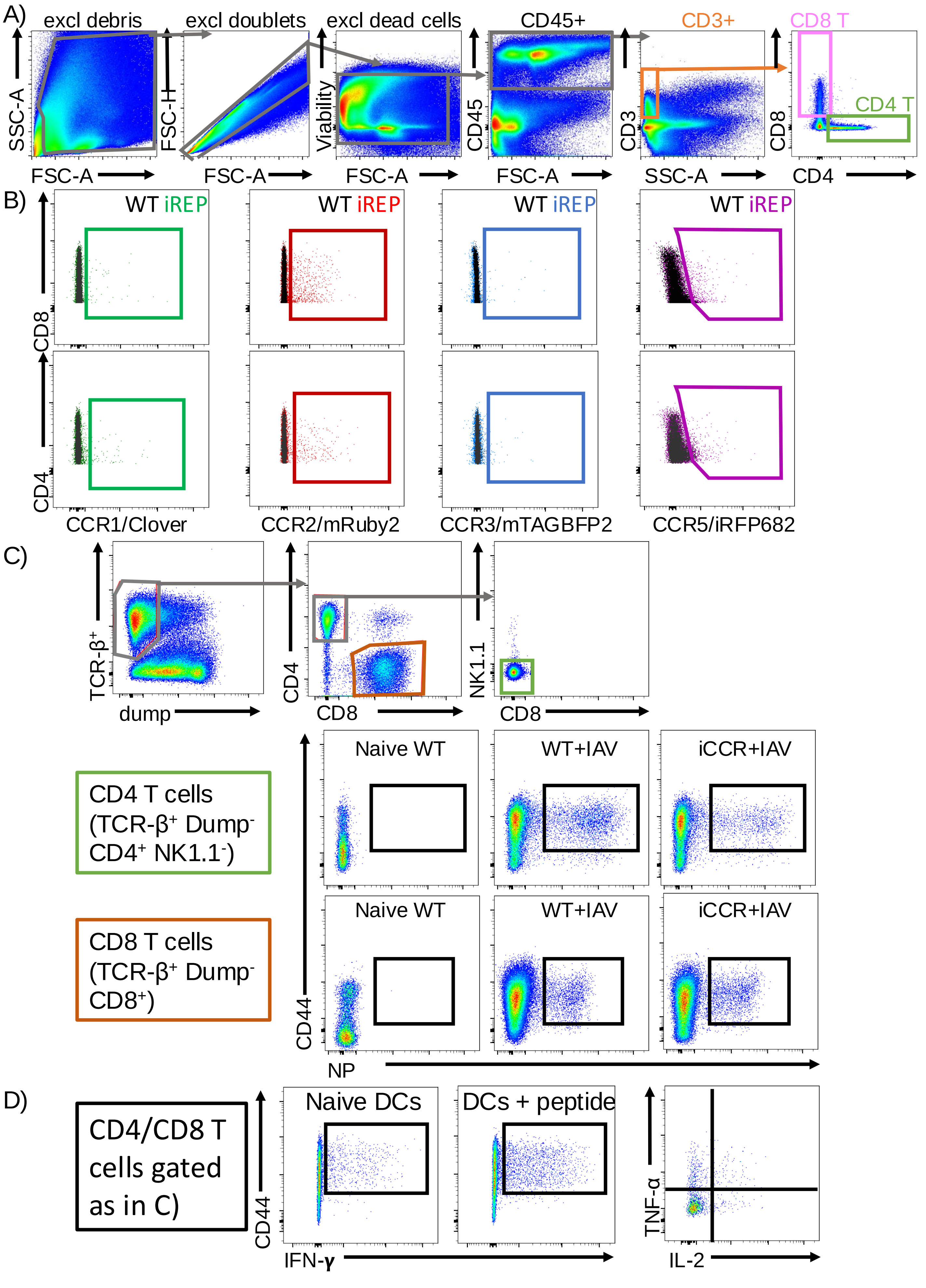

### Supplementary Figure 5. Quantification of T cells and APC in lung and mLN

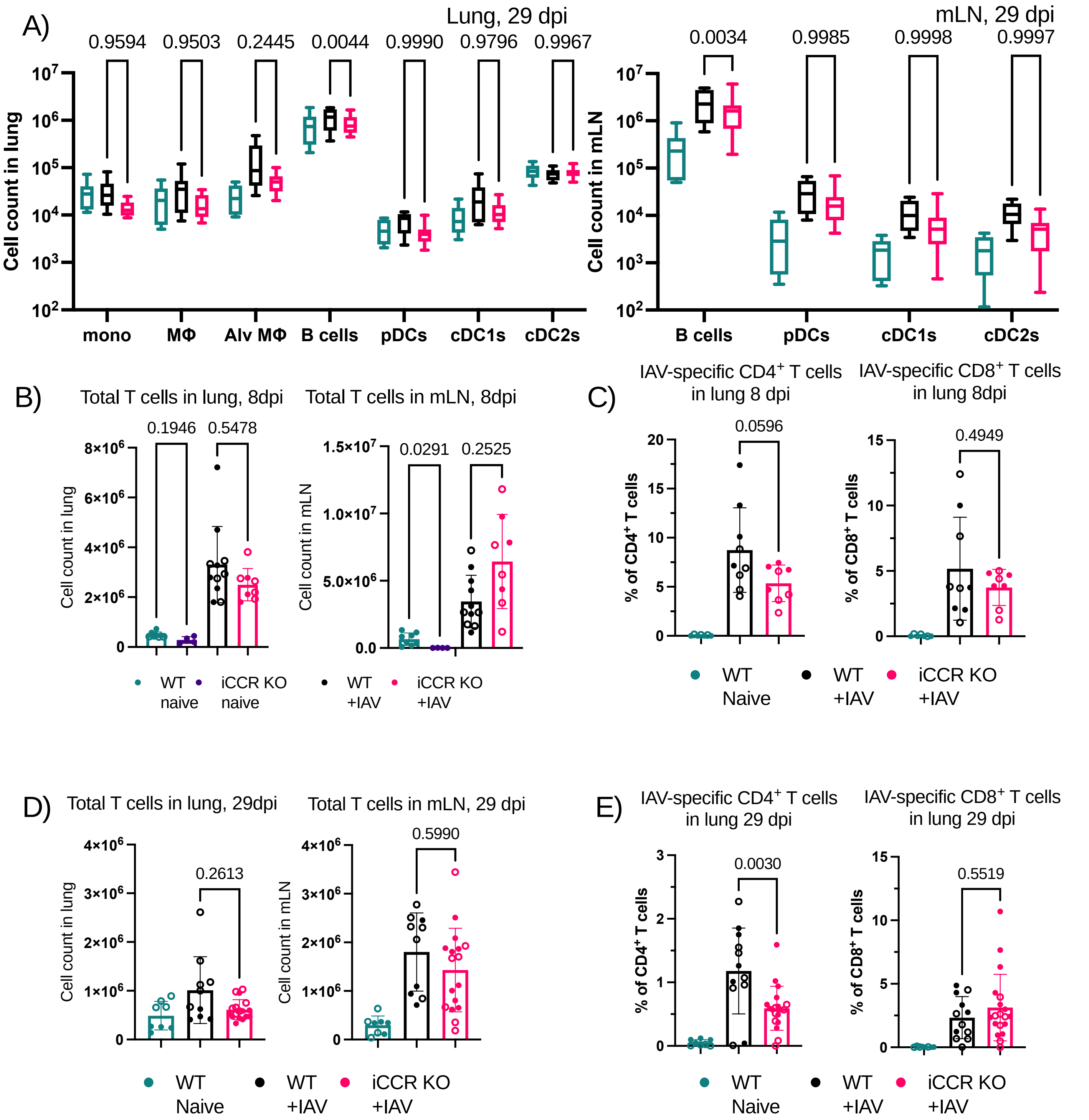
